# Evolution of protection after maternal immunization for respiratory syncytial virus in cotton rats

**DOI:** 10.1101/2021.07.30.454440

**Authors:** Jorge C.G. Blanco, Lori McGinnes-Cullen, Arash Kamali, Fatoumata Y. D. Sylla, Marina S. Boukhavalova, Trudy G. Morrison

## Abstract

Maternal anti-respiratory syncytial virus (RSV) antibodies acquired by the fetus through the placenta protect neonates from RSV disease through the first weeks of life. In the cotton rat model of RSV infections, we previously reported that immunization of dams during pregnancy with virus-like particles assembled with mutation stabilized pre-fusion F protein as well as the wild type G protein resulted in robust protection of their offspring from RSV challenge (Blanco, et al Journal of Virology 93: e00914-19, https://doi.org/10.1128/JVI.00914-19). Here we describe the durability of those protective responses in dams, the durability of protection in offspring, and the transfer of that protection to offspring of two consecutive pregnancies without a second boost immunization. We report that four weeks after birth, offspring of the first pregnancy were significantly protected from RSV replication in both lungs and nasal tissues after RSV challenge, but protection was reduced in pups at 6 weeks after birth. However, the overall protection of offspring of the second pregnancy was considerably reduced, even at four weeks of age. This drop in protection occurred even though the levels of total anti-pre-F IgG and neutralizing antibody titers in dams remained at similar, high levels before and after the second pregnancy. The results are consistent with an evolution of antibody properties in dams to populations less efficiently transferred to offspring or the less efficient transfer of antibodies in elderly dams.

**Author Summary:** Respiratory syncytial virus (RSV) is a major cause of acute lower respiratory tract infection of infants. Because there is no licensed vaccine for RSV as well as potential safety issues with any new vaccine, protection of infants from RSV is problematic. A possible safe approach for infant protection is the transfer of maternal anti-RSV antibodies, induced by immunization, across the placenta to the fetus serving to protect the newborn for months after birth. In a cotton rat model, we have previously shown that maternal immunization with virus-like particles assembled with the RSV F and G proteins protects offspring from RSV infection. Here we describe protection of offspring, following a single immunization, through two pregnancies showing that offspring of the first were well protected from RSV challenge. However, offspring of the second pregnancy were very weakly protected although the levels of total anti-pre-F antibodies and neutralizing antibody titers in the dams remained at constant and high levels before and after the second pregnancy. This result is consistent with an evolution of antibody properties in the dams to those less efficiently transferred to offspring and highlights the importance of appropriate strategies for maternal immunization, such as immunization during each pregnancy.

## Introduction

Respiratory syncytial virus (RSV) is a very common cause of severe acute lower respiratory tract infections in infants and young children, infections that frequently result in hospitalization and, in developing countries, significant mortality (1-3). RSV accounts for approximately three million infections per year world-wide with nearly 200,000 deaths. In the US, RSV infections are the most common cause of infant doctor visits as well as a significant number of hospitalizations (3). However, despite decades of effort, no vaccine has yet been licensed.

Attempts to develop RSV vaccines have been ongoing since the 1960s. Vaccine development has focused on the RSV F protein which is more conserved across all strains of RSV than the G protein and thus should induce protective responses across all strains. The early failures to identify an effective vaccine were due, in part, to a lack of recognition that the pre-fusion conformation of the RSV F protein induces optimal protective responses and that this form of F protein is unstable (4, 5). Another issue is a significant concern about vaccine safety of all candidates stemming from the failure of formaldehyde treated virus (FI-RSV), an early vaccine candidate. FI-RSV was not only ineffective in preventing disease but, more importantly, resulted in life-threatening enhanced respiratory disease (ERD) upon subsequent exposure to infectious RSV (6-9).

Given these considerations, as well as the immunological immaturity of infants and potential interference of maternal antibodies in infant vaccination (10, 11), a current view is that maternal immunization with vaccines containing the mutation stabilized pre-fusion F protein is a safe and efficacious approach for protection of infants against RSV (12-18). Maternal vaccines are commonly used to protect infants from influenza, tetanus, and pertussis (19-22). It has been reported that RSV maternal antibody (matAb) acquired by the fetus through the placenta or lactation has a protective effect on neonates during the first few weeks of life (13, 17, 23-25). Thus, a goal of the maternal RSV immunization strategy is to increase protective maternal antibodies (matAbs) in neonates to levels that will extend the time of their protection after birth.

We have developed novel virus-like particle (VLP) vaccine candidates for RSV. VLPs robustly stimulate immune responses without the complications of adjuvant addition due to their display of repetitive arrays of antigen in a virus-sized particle (26, 27). Because production of VLPs does not require viral replication, multiple antigens and different conformational forms of antigens can be assembled into VLPs, in contrast to attenuated viruses which must remain infectious. VLPs are safer as vaccines for many populations, such as the very young or the very old, compared to infectious, attenuated, or vector viruses since they do not contain a genome and do not produce a spreading infection.

McLellan et al identified mutations in the RSV F protein (DS-Cav1 mutant) that stabilize the pre-fusion conformation, and this prototype pre-fusion F protein is now widely used in many vaccine candidates in different stages of development (5). Subsequently, others have reported different F protein mutations that are stabilizing (28), one of which, UC-3 F, induces higher neutralizing antibody (NAb) titers than the widely-used form, DS-Cav1 F (28, 29). We have assembled VLPs containing the DS-Cav1 or the UC-3 F pre-fusion RSV F proteins along with the RSV G attachment protein and have established UC-3 F VLP superiority over DS Cav1 F as well as post-F protein, or wildtype F protein containing VLPs in inducing NAbs in both mice and cotton rats (28, 30). Furthermore, we have tested the efficacy of these pre-F containing VLPs in the cotton rat model of maternal immunization comparing responses to a post fusion F VLPs as well as soluble versions of the pre- and post-fusion F proteins (18). In these studies, we have used RSV-primed animals in order to mimic normal human populations, most of which have been previously infected with RSV. We have reported that immunization of RSV-primed dams during pregnancy with pre-F VLPs resulted in robust protection of offspring from RSV challenge with no evidence of pathology (29-31). Here we describe the durability of these protective responses in dams after a single immunization and compare the degree of protection these dams can transfer to their offspring in two consecutive litters. We also assess the durability of protection in offspring from each of these two consecutive breedings at extended times after birth.

## Results

### Infection and Immunization

To evaluate the durability of maternal antibody with time after VLP immunization and the transfer of protection to offspring, three-week-old female cotton rats were first infected intranasally (IN) with RSV (RSV-primed, day 0) in order to mimic the RSV-immune condition of the majority of the adult human population. These animals were then set in breeding pairs at 56 days (breed 1) and immunized at two weeks of gestation (day 70) with one of two different pre-fusion F containing VLPs, DS-Cav1 F or UC-3 F (Figure 1). Groups of animals were immunized with 25, 75, 100, or 150 micrograms of UC-3 F VLPs, or 100 micrograms of DS-Cav-1 F VLPs. Control groups were immunized with 100 micrograms of stabilized post fusion F VLPs or re-infected (IN) with RSV (RSV/RSV). Another group of RSV-primed animals did not receive immunization (RSV/mock). Litters of pups delivered after the first breeding on or about (day 84) were divided into two groups, one of which was RSV challenged at 4 weeks after birth while the other was RSV challenged at 6 weeks after birth (Figure 1). RSV-challenged pups were sacrificed four days after the challenge for assessment of serum antibody titers as well as virus titers in lungs and nasal tissue.

**Legend to Figure 1:**
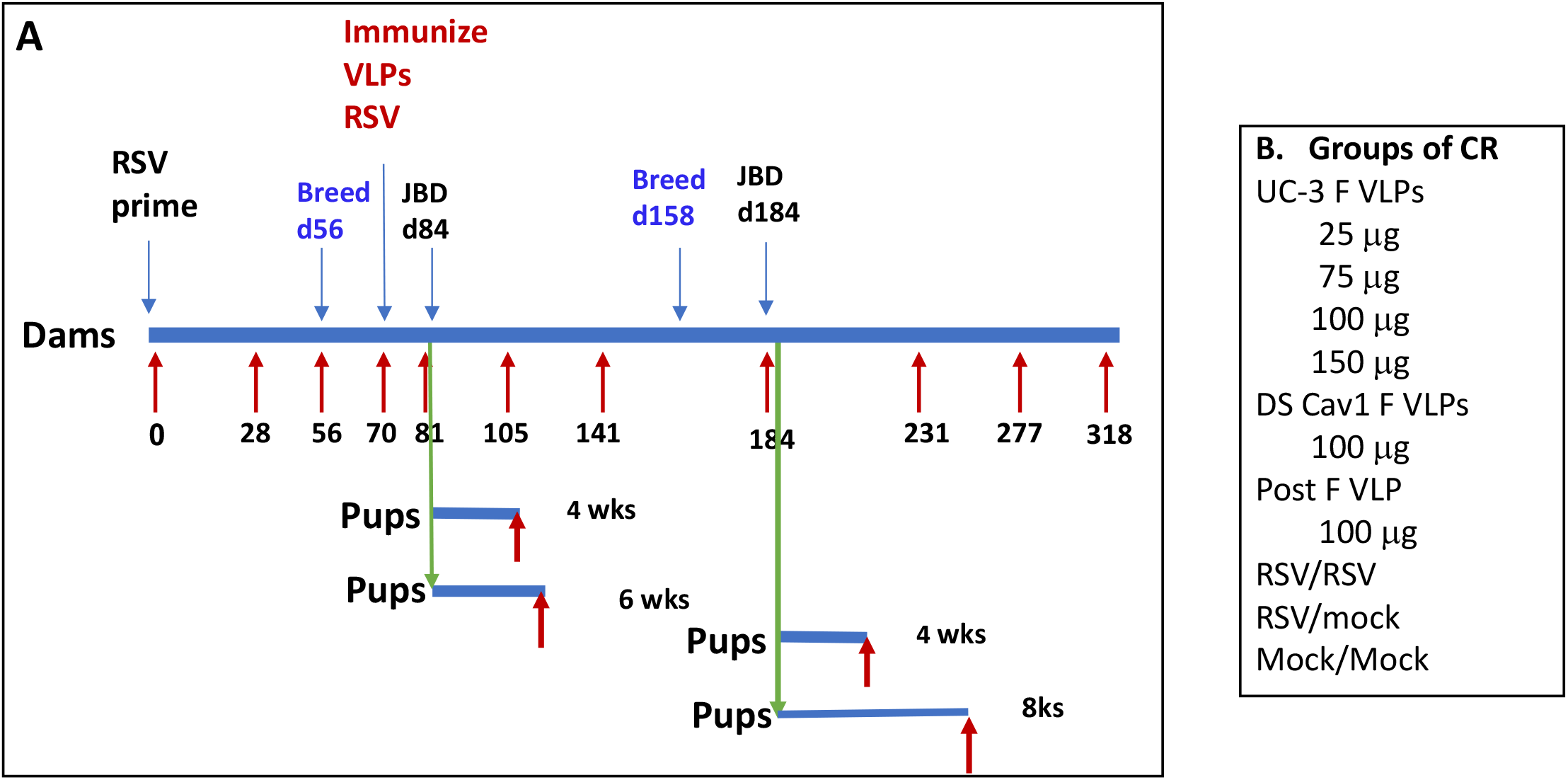
Experimental design. Panel A shows a diagram of the protocol to assess durability of maternal immunization. Groups of CR were primed with RSV (IN) at day 0. These animals were then bred twice, once at day 56, and the second at day 158. Groups of animals were immunized at day 70 with UC-3 F, DS Cav1 F, or post F VLPs. Other groups were infected a second time with RSV or mock immunized. Offspring of dams were challenged with RSV at 4, 6, or 8 weeks after birth. Red arrows show times of serum acquisition. Panel B lists groups of dams (10 CR/group).

The same dam cohort was bred again (breed 2) at 158 days after the RSV prime but without a second immunization (Figure 1). Litters of pups delivered after this second breeding (on or about day 185), were again divided into two groups, one of which was RSV challenged at 4 weeks after birth while the second at 8 weeks. Four days after challenge, virus titers in lungs and nasal tissue were determined. Dams were maintained for a total of 318 days after RSV priming to assess serum antibody levels late in their life. Serum samples from the dams and their offspring were acquired throughout the protocol at intervals indicated by red arrows in Figure 1 for assessment of immune responses with time after immunization.

### Durability of anti-pre-F IgG in dams

Sera from each group of dams from each time point were pooled for determination of the total anti-pre-F IgG by ELISA. The levels of total anti-pre-F IgG antibodies increased significantly after a pre-F VLP or Post F VLP single immunization at day 70, as we have previously reported (18, 28, 31) (Figure 2, panels A and B) and remained relatively stable from day 141 to day 277 varying no more than 20%, with the exception of a drop at day 184 to between 43-58% of that on day 141 (Figure 2, Table 1). Day 184 was just before the delivery of the second breeding. This drop may be due to the transfer of matAb to the fetus just before delivery, a phenomenon we have previously observed in primed, unvaccinated females during the first breeding but not in animals vaccinated during their pregnancy (18). In animals immunized during pregnancy, the serum levels at day 84 are a combination of increasing antibodies due to vaccination and decreasing levels due to their transfer to the fetus. This point is supported by the observation that mock immunized animals were the only group with a decrease in total Pre-F IgG on day 84 (Figure 2, A and B). However, the levels of total pre-F IgG in the dams recovered by day 231 to levels similar to that seen in day 141 (Table 1), suggesting their replenishment after pup delivery by anti-pre-F secreting, bone marrow-associated long-lived plasma cells (LLPC). Indeed, there were increased anti-pre-F IgG secreting LLPC in immunized animals compared to mock immunized animals upon sacrifice of the dams on day 318 (Figure 3).

**Legend to Table 1:** Total ng/ml anti-pre-F IgG in dam sera. The mean ng/ml of anti-Pre-F I IgG in pools of sera from each group of dams from each time point after RSV priming was measured by ELISA using UC-3 F as target. Data from animals immunized with 100 Δg VLPs as well as RSV/RSV and RSV/Mock groups are shown. Numbers, +/-, below each value are the standard deviation of three separate determinations. Numbers in parentheses are the percent of IgG relative to levels at day 141.

| Day Post<br>RSV Prime | Immunogen |  |  |  |  |
| --- | --- | --- | --- | --- | --- |
|  | UC-3F VLP(100ug) | DS Cav1 VLP (100ug) | post F VLP(100 ug) | RSV/RSV | RSV/mock |
| 28 | 1.8x10<sup>5</sup><br>+/-6.9x10<sup>4</sup> | 1.41x10<sup>5</sup><br>+/-4.9x10<sup>4</sup> | 1.10x10<sup>5</sup><br>+/-7.1x10<sup>4</sup> | 1.63x10<sup>5</sup><br>+/-2.6x10<sup>4</sup> | 2.0x10<sup>5</sup><br>+/-4.4x10<sup>4</sup> |
| 56 | 3.5x10<sup>5</sup><br>+/- 1.1x10<sup>5</sup> | 1.92x10<sup>5</sup><br>+/-8.9x10<sup>4</sup> | 2.00x10<sup>5</sup><br>+/-9.2x10<sup>4</sup> | 1.61x10<sup>5</sup><br>+/-5.0x10<sup>4</sup> | 3.3x10<sup>5</sup><br>+/-9.8x10<sup>4</sup> |
| 70 | 3.5x10<sup>5</sup><br>+/-9.7x10<sup>4</sup> | 2.28x10<sup>5</sup><br>+/-9.3x10<sup>4</sup> | 2.66x10<sup>5</sup><br>+/-1.1x10<sup>5</sup> | 3.25x10<sup>5</sup><br>+/-9.2x10<sup>4</sup> | 3.81x10<sup>5</sup><br>+/-1.3x10<sup>5</sup> |
| 84 | 1.19x10<sup>6</sup> (74%)<br>+/-1.7x10<sup>5</sup> | 5.76x10<sup>5</sup> (52%)<br>+/-2.7x10<sup>5</sup> | 5.96x10<sup>5</sup> (87%)<br>+/-2.6x10<sup>5</sup> | 5.66x10<sup>5</sup> (59%)<br>+/-2.1x10<sup>5</sup> | 3.03x10<sup>5</sup> (53%)<br>+/-1.3x10<sup>5</sup> |
| 141 | 1.61x10<sup>6</sup> (100%)<br>+/-4.4x 10<sup>5</sup> | 1.11x10<sup>6</sup> (100%)<br>+/-6.8x10<sup>5</sup> | 6.82x10<sup>5</sup> (100%)<br>+/-2.92x10<sup>5</sup> | 9.58x10<sup>5</sup> (100%)<br>+/-3.6x10<sup>5</sup> | 5.69x10<sup>5</sup> (100%)<br>+/-2.9x10<sup>5</sup> |
| 184 | 6.95x10<sup>5</sup> (43%)<br>+/-1.53x10<sup>5</sup> | 4.93x10<sup>5</sup> (44%)<br>+/-1.6x10<sup>5</sup> | 3.94x10<sup>5</sup> (58%)<br>+/-1.0x10<sup>5</sup> | 4.01x10<sup>5</sup> (41%)<br>+/-8.8x10<sup>4</sup> | 3.89x10<sup>5</sup> (68%)<br>+/-2.1x10<sup>5</sup> |
| 231 | 1.80 x10<sup>6</sup> (112%)<br>+/-7.13x10<sup>5</sup> | 9.35x10<sup>5</sup> (84%)<br>+/-3.7x10<sup>5</sup> | 7.85x10<sup>5</sup> (115%)<br>+/-1.9x10<sup>5</sup> | 1.01x10<sup>6</sup> (105%)<br>+/-3.2x10<sup>5</sup> | 6.55x10<sup>5</sup> (115%)<br>+/-2.8x10<sup>5</sup> |
| 277 | 1.58x10<sup>6</sup> (98%)<br>+/-6.21x10<sup>5</sup> | 9.13x10<sup>5</sup> (82%)<br>+/-2.9x10<sup>5</sup> | 7.51x10<sup>5</sup> (110%)<br>+/-4.9x10<sup>4</sup> | 7.73x10<sup>5</sup> (80%)<br>+/-2.1x10<sup>5</sup> | 6.23x10<sup>5</sup> (100%)<br>+/-2.2x10<sup>5</sup> |
| 318 | 1.30x10<sup>6</sup> (81%)<br>+/-2.66x10<sup>5</sup> | 6.10x10<sup>5</sup> (55%)<br>+/-3.7x10<sup>4</sup> | 7.39x10<sup>5</sup> (108%)<br>+/-3.1x10<sup>5</sup> | 5.67x10<sup>5</sup> (59%)<br>+/-2.9x10<sup>4</sup> | 5.58x10<sup>5</sup> (98%)<br>+/-2.6x10<sup>5</sup> |

**Legend to Figure 2:**
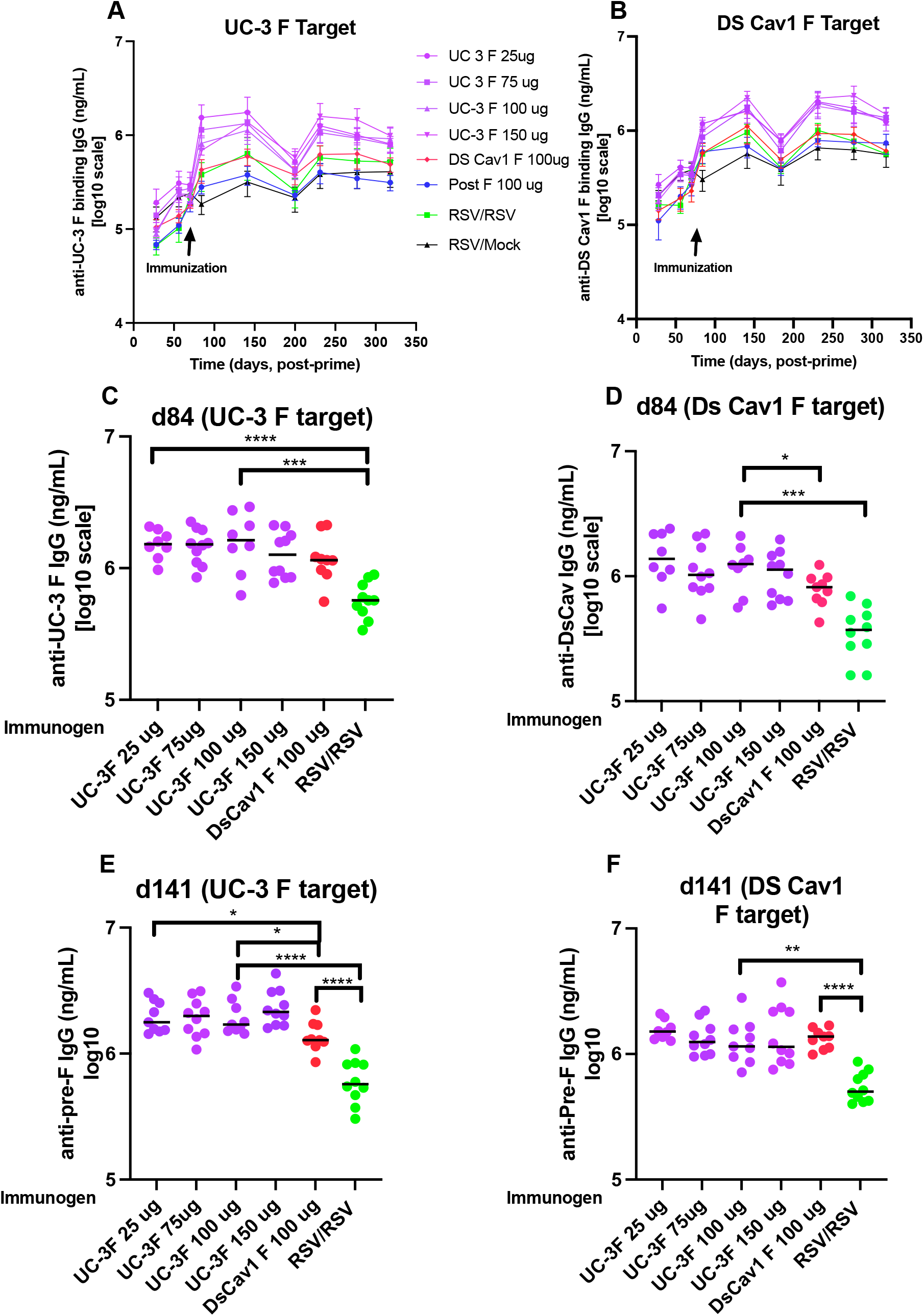
Total ng/ml of serum anti-pre-F protein IgG. The concentrations of anti-pre-F serum IgG in different groups of dams immunized with VLPs or RSV were assessed by ELISA using soluble UC-3 F (panels A, C, E) or soluble DS-Cav1 F (panels B, D, F) as target. Serum samples acquired in different groups of animals immunized with VLPs at each time point were pooled (panels A, B) and error bars show mean and standard deviation of three separate determinations. Results from serum in individual animals immunized with different concentrations of UC-3 F VLPs or 100 Δg DS Cav1 F VLPs and acquired at days 84 or 141 are shown in panels C-F. Mean for each group is shown as a horizontal black line. *p<0.05; **p<0.01; ***p<0.001; ****p<0.0001

**Legend to Figure 3:**
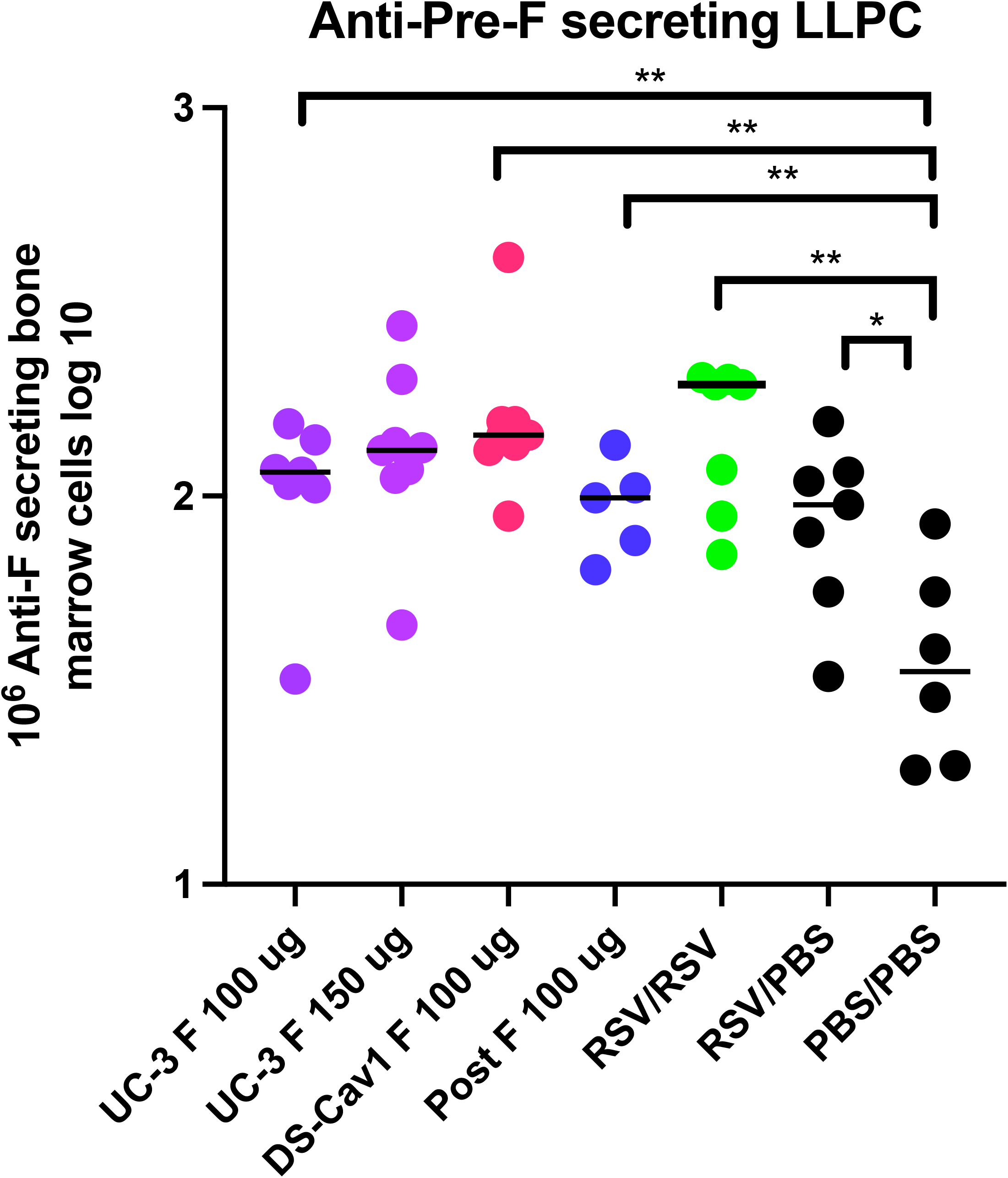
Bone marrow associated long-lived plasma cells (LLPC) secreting anti-pre-F IgG. At day 318, bone marrows of dams were acquired and the numbers of cells secreting anti-pre-F IgG were measured as described in Methods. Results from bone marrows of individual animals are shown as log_10_ of the numbers of positive cells in 10^6^ cells. Immunogen in dams is shown on the x axis. Solid line indicates mean titers in each data set. *p<0.05; **p<0.01.

Measured levels of total anti-pre-F IgG in all serum samples were consistently slightly higher when soluble UC-3 F instead of soluble DS-Cav1 was used as target for ELISA (Figure 2, panels A and B) suggesting either that the UC-3 F target detects a broader range of antibody specificities than the DS-Cav1 target or that the affinity of antibodies to UC-3 F target is greater than to DS-Cav1. At most time points and with either target, levels of total anti-pre-F IgG were higher in sera of UC-3 F VLP immunized animals compared to DS-Cav1 F VLP immunized animals (Figure 2, A, B). Titers induced by either pre-fusion F VLPs were higher than that induced by post F VLPs or two consecutive RSV infections (Figure 2). Immunization with different amounts of UC-3 F VLPs made little difference in total anti-pre-F IgG at all times, illustrated by anti-pre-F IgG in sera from individual animals at days 84 and 141 (Figure 2, panels C-F) (Table 2).

**Legend to Table 2:** Effect of different doses of UC-3 F VLPs on total sera ng/ml anti-pre-F IgG. The mean ng/ml of anti-pre-F IgG in pools of sera from each of four groups of UC-3 F VLP immunized dams at each time point after RSV priming was measured in ELISA using UC-3 F as target. Numbers, +/-, below each value are the standard deviations of three separate determinations. Numbers in parentheses are the percent of IgG relative to levels at day 141.

| Day Post<br>RSV Prime | Immunogen |  |  |  |
| --- | --- | --- | --- | --- |
|  | UC-3F VLP(25 µg) | UC-3 F VLP (75µg) | UC-3 F (100 µg) | UC-3 F (150 µg) |
| 28 | 2.68x10<sup>5</sup><br>+/-1.2810<sup>5</sup> | 2.07x10<sup>5</sup><br>+/-6.9x10<sup>4</sup> | 1.81x10<sup>5</sup><br>+/-6.91x10<sup>4</sup> | 2.06x10<sup>5</sup><br>+/-9.49x10<sup>4</sup> |
| 56 | 4.09x10<sup>5</sup><br>+/- 1.33x10<sup>5</sup> | 3.43x10<sup>5</sup><br>+/-7.74x10<sup>4</sup> | 3.52x10<sup>5</sup><br>+/-1.13x10<sup>5</sup> | 3.7x10<sup>5</sup><br>+/-7.61x10<sup>4</sup> |
| 70 | 4.04x10<sup>5</sup><br>+/-1.14x10<sup>5</sup> | 3.34x10<sup>5</sup><br>+/-5.98x10<sup>4</sup> | 3.57x10<sup>5</sup><br>+/-9.77x10<sup>4</sup> | 3.40x10<sup>5</sup><br>+/-1.16x10<sup>5</sup> |
| 84 | 1.18x10<sup>6</sup> (72%)<br>+/-3.41x10<sup>5</sup> | 8.53x10<sup>5</sup> (48%)<br>+/-2.99x10<sup>5</sup> | 1.19x10<sup>6</sup> (74%)<br>+/-1.76x10<sup>5</sup> | 9.87x10<sup>5</sup> (44%)<br>+/-1.00x10<sup>5</sup> |
| 141 | 1.64x10<sup>6</sup> (100%)<br>+/-7.53x 10<sup>5</sup> | 1.77x10<sup>6</sup> (100%)<br>+/-7.32x10<sup>5</sup> | 1.61x10<sup>6</sup> (100%)<br>+/-4.44x10<sup>5</sup> | 2.24x10<sup>6</sup> (100%)<br>+/-6.65x10<sup>5</sup> |
| 184 | 7.84x10<sup>5</sup> (48%)<br>+/-2.50x10<sup>5</sup> | 6.12x10<sup>5</sup> (35%)<br>+/-2.52x10<sup>5</sup> | 7.75x10<sup>5</sup> (48%)<br>+/-9.4x10<sup>4</sup> | 7.90x10<sup>5</sup> (35%)<br>+/-1.91x10<sup>5</sup> |
| 231 | 2.03x10<sup>6</sup> (126%)<br>+/-8.59x10<sup>5</sup> | 1.98x10<sup>6</sup> (112%)<br>1.10x10<sup>5</sup> | 1.81x10<sup>6</sup> (112%)<br>+/-7.13x10<sup>5</sup> | 2.21x10<sup>6</sup> (98%)<br>+/-5.75x10<sup>5</sup> |
| 277 | 1.73x10<sup>6</sup> (108%)<br>+/-6.21x10<sup>5</sup> | 1.58x10<sup>6</sup> (89%)<br>+/-2.9x10<sup>5</sup> | 1.58 x10<sup>6</sup> (98%)<br>+/-4.9x10<sup>4</sup> | 2.36x10<sup>6</sup> (105%)<br>+/-2.1x10<sup>5</sup> |
| 318 | 1.24x10<sup>6</sup> (73%)<br>+/-1.50x10<sup>5</sup> | 1.38x10<sup>6</sup> (78%)<br>+/-6.68x10<sup>5</sup> | 1.30x10<sup>6</sup> (81%)<br>+/-2.66x10<sup>5</sup> | 1.49x10<sup>6</sup> (66%)<br>+/-4.14x10<sup>5</sup> |

### Durability of neutralizing antibody (NAb) in dams

Serum NAb titers stimulated in dams by UC-3 F VLPs were higher than titers stimulated by DS Cav1 VLPs (Figure 4 A, B), as we have previously reported (28), and titers after either pre-F VLP immunization were higher than after post-F VLP immunization (Figure 4, panels A-C). Similar to levels of total anti-pre-F IgG, the dose of UC-3 F VLPs had little impact on NAb titers at all times after immunization (Figure 4, Panels F-H).

**Legend to Figure 4:**
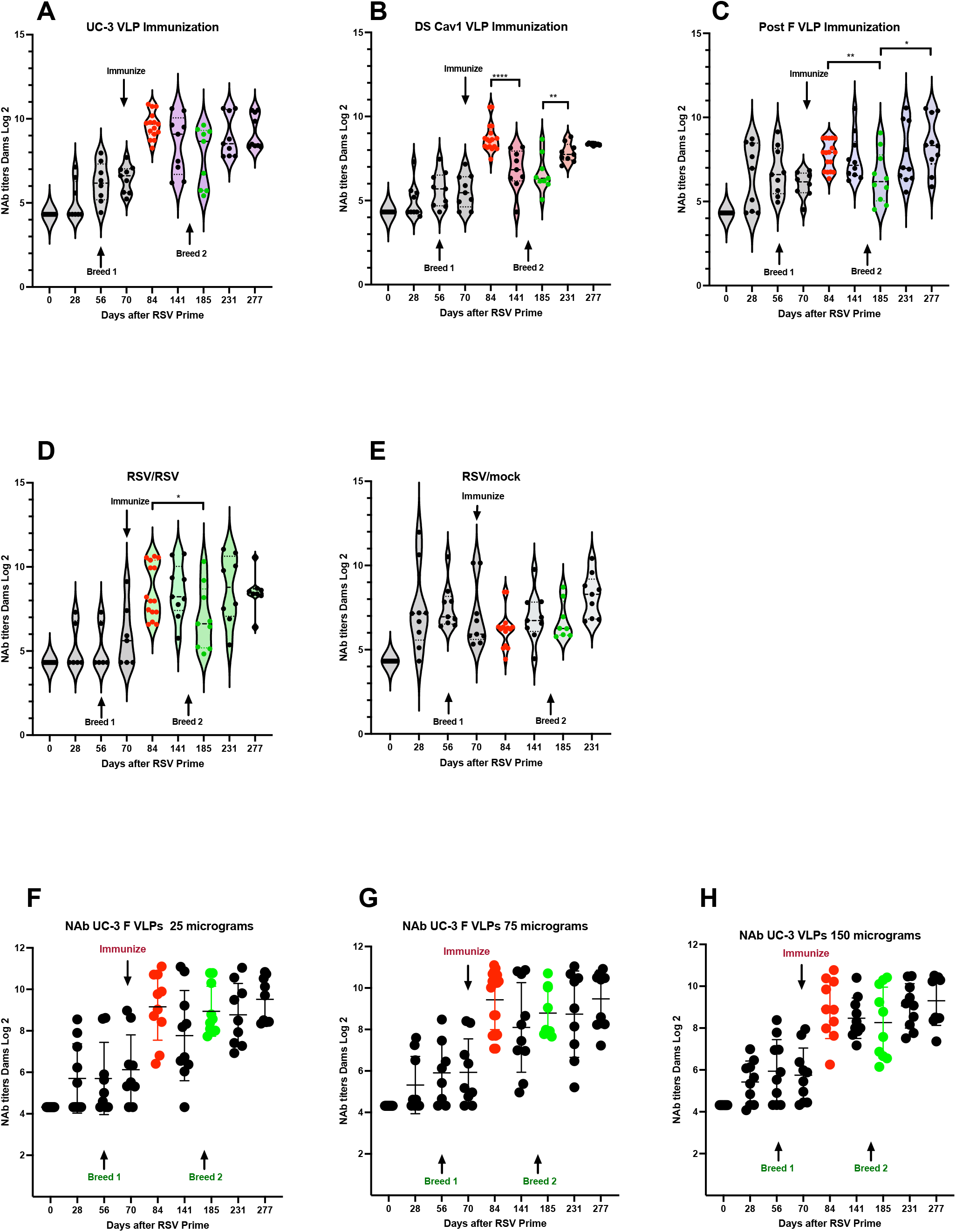
Neutralization titers in dams. The serum NAb titers in dams, primed with RSV at day 0 and immunized at day 70, at each time point are shown. NAb titers of sera from animals immunized with 100 μg VLP protein are shown in panels A-C while titers in animals immunized with RSV or mock immunized are shown in panels D and E. Panels A-E are presented as violin plots with individual animal data shown. Mean and standard deviation are indicated by solid and dotted black lines, respectively. Panels F-H show results of individual animals immunized with 25, 75, or 150 μg of UC-3 F VLPs. Mean of each group is shown as a black horizontal bar. Red and green dots: data from sera acquired just before delivery of breed 1 or 2, respectively. *p<0.05; **p<0.01; ***p<0.001; ****p<0.0001

In contrast to total anti-pre-F IgG, NAb titers in the dams after immunization with UC-3 F VLPs remained relatively stable throughout the time course with no statistical differences between time points of day 84 to 277 (Figure 4). However, titers after DS-Cav1 VLP immunization dropped at day 141 and 184, and then recovered by day 231 (Figure 4A and B). Titers after post F VLP immunization dropped slightly at day 184 and then recovered by day 277. Thus, the levels of dams’ NAb titers after pre-F VLP immunization did not necessarily track with the titers of total anti-pre-F IgG in the respective animals and levels varied with the immunogen. These results suggested that the NAbs generated in the primed dams were a subpopulation of the total anti-pre-fusion F IgG induced by immunization.

### Comparison of total serum anti-pre-F IgG in breed 1 and breed 2 pups

Since levels of total anti-pre-F IgG in the sera of dams remained relatively constant from day 84 to day 277 with the exception of a transient drop at day 184 (which may reflect transfer to offspring, Figure 2A and B), we asked if the levels of total maternal pre-F IgG (matAbs) transferred to offspring were similar in breed 1 vs breed 2. Anti-pre-F IgG levels in breed 1 offspring of pre-F VLP immunized dams or RSV immunized dams at four weeks after birth were on average 15 to 20% that in the dam sera at day 141 (Figure 5E, F) (Table 3) while that in offspring of dams immunized with post-F VLP was 35% that in dams. However, in breed 2 pups, the total anti-pre-F IgG was, on average, 6 to 11% that in dams at day 141 and 30-53% that in sera of breed 1 pups (Figure 5, panels E and F) (Table 3) although dam anti-pre-F IgG did not vary significantly between days 141 and 231. These results suggest that the population of anti-pre-F IgG antibodies in immunized dams is less efficiently transferred to pups in breed 2 compared to breed 1. As in the dam sera, the concentration of UC-3 F VLP used as immunogen had little effect on the pup IgG titers (Figure 5, panels A-D).

**Legend to Table 3:** Comparisons of ng/ml off sera anti-pre-F IgG in dams and pups. The mean and standard deviations, respectively, of titers of anti-pre-F IgG, measured by ELISA, in each of the 8 groups of dams at day 141 is shown on the first two lines. Lines 3 and 4 show means and standard deviations, respectively, of anti-pre-F IgG titers in sera of breed 1 offspring of each of the 8 groups of dams at four weeks after birth while lines 5 and 6 show similar data for offspring of breed 2. Line 7 shows breed 1 pup titers as a percent of dam titers at day 141 while line 8 shows breed 2 pup titers as a percent of dam titers at day 141.

| Total<br>Pre-F IgG | Dams Immunization |  |  |  |  |  |  |  |
| --- | --- | --- | --- | --- | --- | --- | --- | --- |
|  | UC-3 25 | UC-3 75 | UC-3 100 | UC-3 150 | DS Cav1 100 | Post F 100 | RSV/RSV | RSV/PBS |
| Dams<br>Day 141 | 1.63x10<sup>6</sup><br>+/-7.5 x 10<sup>5</sup> | 1.77 x 10<sup>6</sup><br>+/-7.3x10<sup>5</sup> | 1.61 x 10<sup>6</sup><br>+/-4.4x10<sup>5</sup> | 1.62x 10<sup>6</sup><br>+/-6.6x10<sup>5</sup> | 1.11x10<sup>6</sup><br>+/-6.7x10<sup>5</sup> | 6.82 x 10<sup>5</sup><br>+/-2.92x10<sup>5</sup> | 9.59 x 10<sup>5</sup><br>+/-3.6x10<sup>5</sup> | 5.69 x 10<sup>5</sup><br>+/-2.9x10<sup>5</sup> |
| Pups<br>Breed 1<br>4 wks | 2.40x10<sup>5</sup><br>+/-6.7x10<sup>4</sup> | 3.55x10<sup>5</sup><br>+/-1.1x10<sup>4</sup> | 3.34x10<sup>5</sup><br>+/-5.6x10<sup>4</sup> | 3.30x10<sup>5</sup><br>+/-5.1x10<sup>4</sup> | 2.27x10<sup>5</sup><br>+/-6.8x10<sup>4</sup> | 2.39x10<sup>5</sup><br>+/-1.8x10<sup>4</sup> | 1.96x10<sup>5</sup><br>+/- 7.0x10<sup>4</sup> | 8.39 x 10<sup>4</sup><br>+/-2.9x10<sup>4</sup> |
| Pups<br>Breed 2<br>4 wks | 1.37x10<sup>5</sup><br>+/-4.0x10<sup>4</sup> | 1.44x10<sup>5</sup><br>+/-6.7x10<sup>4</sup> | 1.24x10<sup>5</sup><br>+/-5.6x10<sup>4</sup> | 1.37 x10<sup>5</sup><br>+/-2.3x10<sup>4</sup> | 1.08x10<sup>5</sup><br>+/-6.8x10<sup>4</sup> | 7.3x10<sup>4</sup><br>+/-3.6x10<sup>4</sup> | 6.5x10<sup>4</sup><br>+/-2.4x10<sup>4</sup> | 5.7x10<sup>4</sup><br>+/-2.1x10<sup>4</sup> |
| % of Dam<br>Breed 1 | 15 | 20 | 21 | 20 | 20 | 35 | 20 | 15 |
| % of Dam<br>Breed 2 | 8 | 8 | 7 | 8 | 10 | 11 | 6 | 10 |

**Legend to Figure 5:**
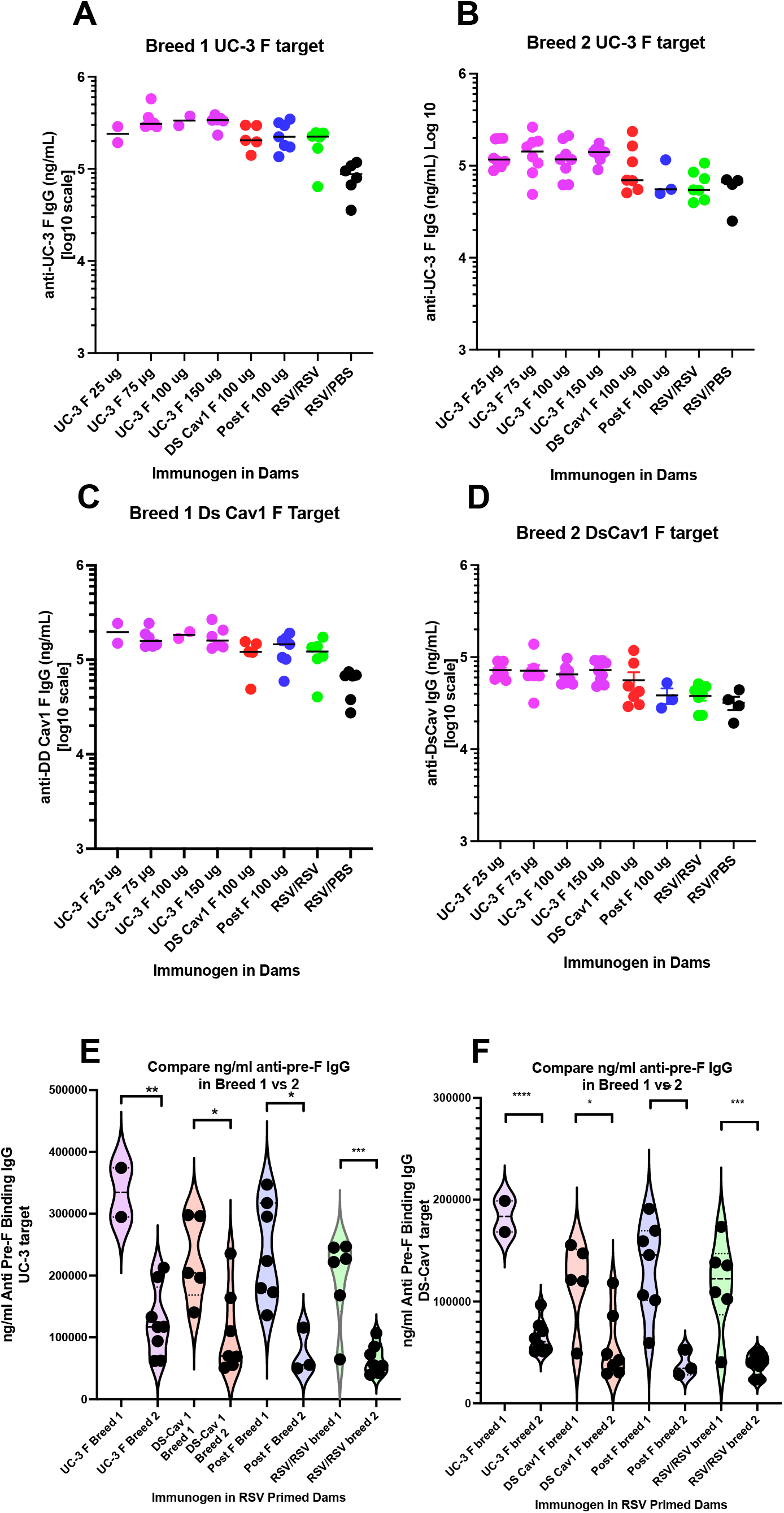
Comparisons of total anti-pre-F IgG in offspring of immunized dams. Shown are total anti-pre-F IgG (ng/ml) (shown on a log_10_ scale) in individual offspring of breed 1 (panels A, and C) and breed 2 (panels B, D) dams immunized with 25, 75, 100, and 150 Δg of UC-3F VLP, 100 Δg of DS-Cav1 VLPs, 100 Δg post F VLPs, RSV, or mock immunized. Pre-F IgG titers at four weeks after birth were determined using as target in ELISA soluble UC-3 F (panels A, B) or DS Cav1 F (panels C, D). Mean is indicated by horizontal black line. Panels E and F directly compare levels of total anti-pre-F IgG in sera from breed 1 and 2 offspring of dams immunized with 100 Δg of VLPs using soluble UC-3 F or DS Cav1 F or as target, respectively, and shown on a linear scale. Mean and standard deviation are indicated by dashed and dotted black line, respectively. *p<0.05; **p<0.01; ***p<0.001; ****p<0.0001.

### Comparisons of matNAb titers in sera of breed 1 and breed 2 pups

The matNAb (maternal neutralizing antibodies) titers in sera of all pups from breed 1 were considerably higher than that in sera of breed 2 pups irrespective of the immunogen used for dams (Figure 6, panels A and B, respectively) and consistent with the differences in total anti-pre-F IgG between these two breeds of pups (Figure 5). In this case the dose of the immunogen in dams made some difference in titers on matNAb in both breeds. The 100 μg dose of UC-3 F VLPs resulted in the highest mean matNAb titers, however, the mean titers in breed 2 pup sera were considerably lower than those in breed 1 sera, with the mean titer in breed 1 at 7, log_2_, and in breed 2 at 5, log_2_. The mean serum matNAb titers in both breeds of offspring of dams immunized with DS-Cav-1 or post F VLPs were considerably lower than titers resulting from immunization with a similar concentration of UC-3 F VLPs.

**Legend to Figure 6:**
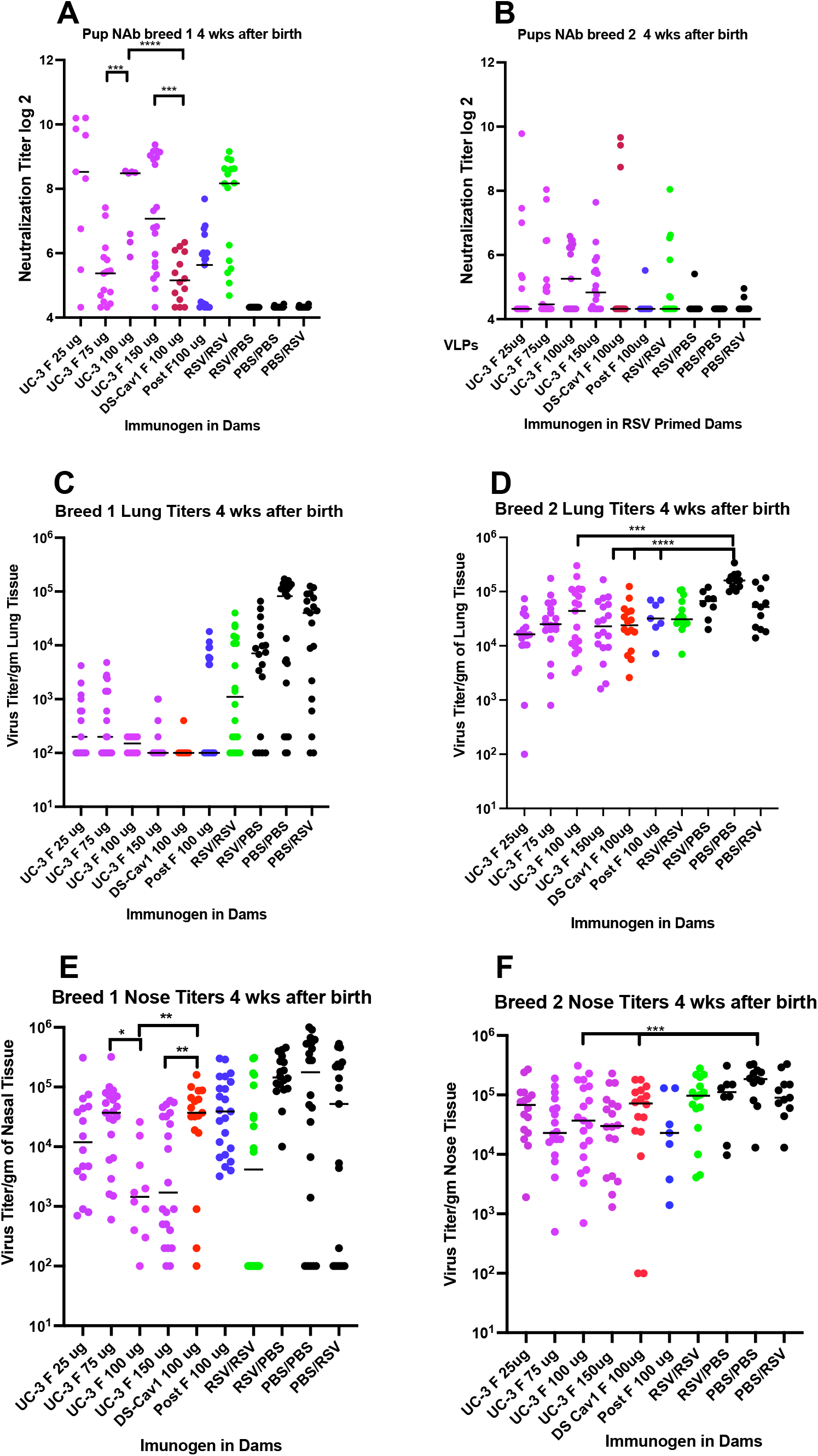
Protective Responses in offspring of immunized dams. Panels A, B: neutralizing antibody titers in sera of individual offspring from breed 1 (panel A) or breed 2 (panel B). Panels C and D: RSV titers in lungs of individual offspring from breed 1 or 2, respectively, after RSV challenge at 4 weeks after birth. Panels E and F: RSV titers in nasal tissue of individual offspring from breed 1 or 2, respectively, after RSV challenge at 4 weeks after RSV challenge. Mean of each data set is indicated by solid black line. *p<0.05; **p<0.01; ***p<0.001; ****p<0.0001

### Comparisons of protection by matAb of breed 1 and breed 2 pups from RSV challenge

Titers of virus in lungs and nasal tissues of pups after RSV challenge directly reflects the extent of protection of the pups by maternal antibodies. The lung and nasal tissue RSV titers in breed 1 pups (Figure 6, panel C and E respectively) clearly demonstrate that all breed 1 offspring of dams immunized with any pre-F VLP were significantly protected from replication of the virus in lung tissue, reducing virus titers by three Log_2_ (panel C). However, only UC-3 F VLP immunization resulted in significant protection from replication in nasal tissues (panel E). Immunization with 100 μg of UC-3 F VLPs resulted in a decrease of two Log_2_ of virus (panel E) in nasal tissue, while maternal immunization using DS-Cav1 F or post-F VLP resulted in, at best, a half log_2_ reduction in nasal tissue titer. This result indicated that UC-3 F VLPs immunization conferred more robust protection than DS Cav1 F VLPs consistent with results previously reported (28).

In contrast, pups born from breed 2 showed much lower protection from replication of the virus in lung tissue and nasal tissue (Figure 6, panels D and F, respectively). At best, there was a one Log_2_ reduction in virus titers in these tissues, a result consistent with the lower levels of NAb in these breed 2 animals. Thus, while total anti-pre-F IgG in dams remained relatively constant, the transfer of matNAbs to their respective pups decreased during a second breeding and correlated with the decrease in protection. This result may reflect an evolution of the properties of dam antibodies, with time after immunization, to populations less efficiently transferred to offspring. Alternatively, placental transfer may not be as efficient in elderly animals.

### Durability of Transferred of total anti-pre-F IgG and NAb to offspring

Since there was significant protection of offspring from RSV challenge four weeks after birth, particularly in breed 1 pups, we determined the durability of this protection by comparing the levels total anti-pre-F IgG antibodies, matNAb, and virus titers in lung and nasal tissue in breed 1 pups at 4 and at 6 weeks after birth. Comparisons of the total anti-pre-F IgG showed levels at 6 weeks of 40 to 61% that at 4 weeks regardless of the dose of antigen in the dams (Figure 7, panels A, B). The mean matNAb titers at 6 weeks after birth were 3 Log_2_ lower than at 4 weeks after birth (Figure 7, panels C-D). These differences in the matNAb titers translated to different levels of protection at 4 and 6 weeks after birth from RSV challenge. After challenge, RSV titers in lungs or in nasal tissue at 4 weeks after birth were 1-2 Log_10_ lower than titers at 6 weeks after birth (Figure 7, panels E-H). Thus, in breed 1 pups, the matNAb titers and levels of protection from an RSV challenge diminished significantly by 6 weeks after birth, however, the matNAb titers in UC-3 VLP-vaccinated animals remain significantly higher and lung and nose RSV titers significantly lower than in control offspring of RSV/mock, RSV/RSV, or unvaccinated dams. Furthermore, the durability of protection of offspring of UC-3 F VLP immunized dams was significantly better than that of offspring of DS Cav1 immunized dams.

**Legend to Figure 7:**
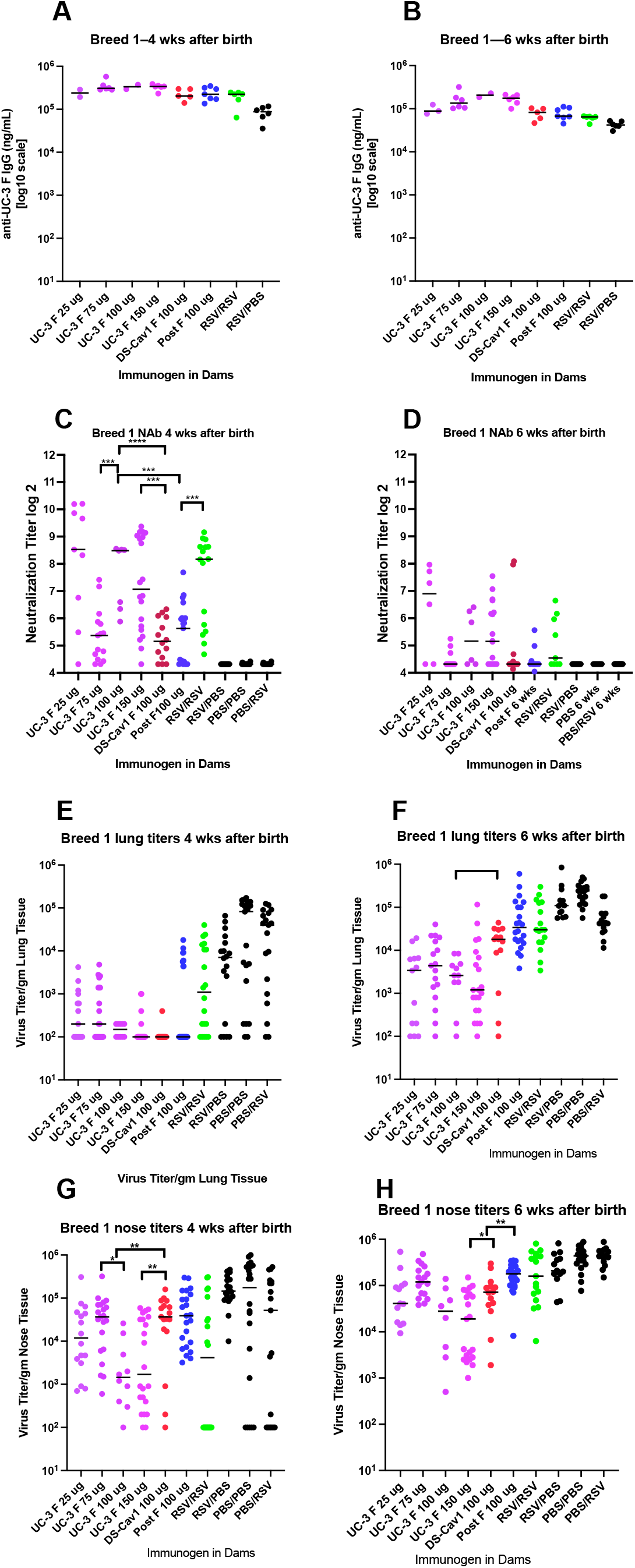
Durability of protection in breed 1 offspring. Sera and tissues acquired at 4 weeks (panels A, C, E, G) or 6 weeks (panels B, D, F, H) after birth of breed 1 pups were assessed for total anti-pre-F IgG (panels A, B), NAb titers (panels C, D), lung titers (panels E, F) or nasal tissue titers (panels G, H) after RSV challenge; *p<0.05; **p<0.01; ***p<0.001; ****p<0.0001.

Breed 2 pups were similarly characterized at 4 weeks and 8 weeks after birth (Figure 8). While levels of total pre-F IgG were surprisingly similar at 4 and 8 weeks (Figure 8, panels A, B) the levels of matNAbs were strikingly different (Figure 8, panels C and D). Breed 2 pups at 8 weeks after birth had no detectable matNAb, and were not protected from virus challenge (Figure 8, panels F, I). Thus, levels of protection from RSV challenge in these breed 2 pups is strongly correlated with the presence of matNAbs, and their steady decrease with the time after birth to undetectable levels by 8 weeks after birth. This loss of protection was despite the presence of significant levels of total anti-pre-F maternal IgG antibodies remaining in these animals.

**Legend to Figure 8:**
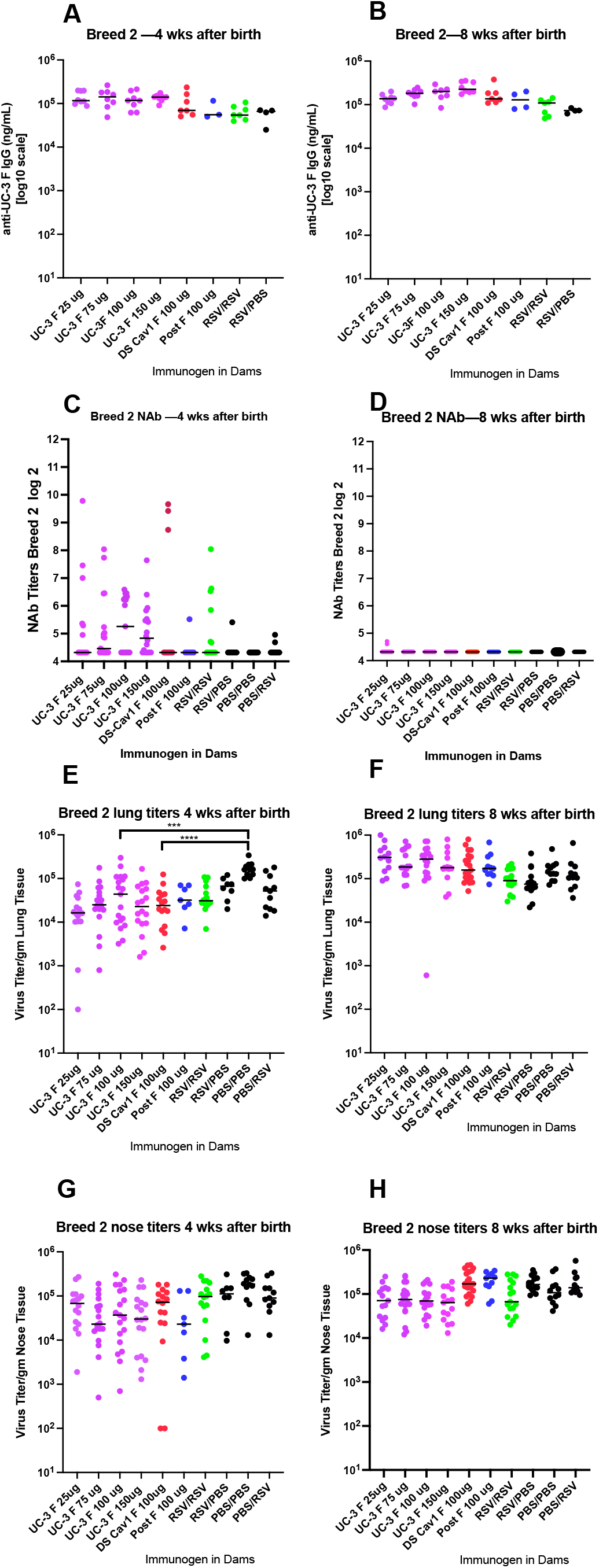
Durability of protection in breed 2 offspring. Sera and tissues acquired at 4 weeks (panels A, C, E, G) or 8 weeks (panels B, D, F, H) after birth of breed 2 pups were assessed for total anti-pre-F IgG (panels A, B), NAb titers (panels C, D), RSV lung titers (panels E, F) or nasal tissue titers (panels G, H) after RSV challenge; ***p<0.001; ****p<0.0001.

## Discussion

Maternal immunization for protection of offspring from some pathogens, including influenza, tetanus, and pertussis infections, is routinely used (19-22)). We have previously reported that, in the cotton rat model of RSV infection, immunization of dams with virus-like particles assembled with mutation stabilized pre-fusion F proteins as well as wild type G protein robustly protected offspring from RSV replication in lungs and nasal tissues after virus challenge (18, 28). Here our goal was to assess the durability of these protective immune responses in dams through two pregnancies and the durability of transfer of protection to offspring of these two breeding events. A secondary goal was to determine the influence of different F protein stabilizing mutations in induction, durability, and transfer of protective responses to offspring as well as the role of different doses of immunogen on levels of these responses.

First, results showed that after a single immunization of RSV primed animals with VLPs during the first pregnancy, the levels of total anti-pre-F IgG in dams remained stable for up to 277-318 days after RSV priming with the exception of a significant but transient drop in those levels just before delivery of offspring of the second breeding, on day 184. This drop in IgG is consistent with the transfer of antibodies to the pups. However, the anti-pre-F IgG levels in dams rebounded to levels before the second pregnancy (day 141) suggesting replenishment by bone marrow associated long-lived plasma cells (LLPC) secreting anti-pre-F IgG. Indeed, these cells were detected in dams on day 318 at the end of the protocol. Thus, levels of these antibodies in immunized dams were maintained for most of the life of the animal.

We also found that offspring protection and matNAb responses induced by UC-3 F VLPs were superior and more durable than those induced by DS Cav1 F VLPs as well as post-F VLPs, confirming our previous reports of the superiority of the UC-3 F VLPs (28). Additional findings reported here are that protective responses in dams did not change significantly or reproducibly with increased doses of immunogen. Furthermore, the results show that immunization with VLPs assembled with F and G proteins resulted in superior protective responses compared to dams infected with one or two doses of RSV, a result that suggests that vaccine candidates may be developed that can provide better protection than a natural RSV infection.

Perhaps the most significant finding in this study was that, while offspring of the first breeding of the immunized dams were robustly protected from RSV challenge, the protection afforded to offspring of the second breeding was considerably reduced. Breed 1 pups had significantly higher levels of anti-pre-F IgG and NAb titers as well as sharply decreased levels of RSV in lung and nasal tissue following RSV challenge compared to breed 2 pups. These findings were surprising since the total anti-pre-F IgG levels in the dams were at high levels prior to the second breeding and there was evidence of the transfer of the antibodies to the pups just before the second delivery. In addition, the levels of NAb, at least in UC-3 F VLP immunized dams, were constant well past the second delivery. The implication of this finding is that dam antibodies were much less efficiently transferred to their offspring in the second breeding. Indeed, the mean of levels of anti-pre-F IgG in breed 2 pups was 30-53 % lower than levels in breed 1 pups. This finding suggests that the antibodies in dams evolved in a way to decrease efficiency of placental antibody transfer. Alternatively, placental transfer of antibodies may be less efficient in elderly animals.

Time post vaccination could result in changes in the composition of the population of MatAbs transferred. This possibility is suggested by the surprising observation that breed 2 pup sera had similar and significant levels of total anti-pre-F IgG antibodies at 4 and 8 weeks but no matNAbs could be measured at 8 weeks. In addition, in cotton rats as in humans, maternal antibodies, mostly IgA, are transferred to the newborn through lactation (32). Our data would then suggest that the protective antibodies transferred through this route may also be reduced by either consecutive pregnancies, time after immunization, and/or the age of the females. Potentially, maternal vaccination against RSV during each pregnancy will be necessary to boost the generation of protective MatNAbs that can be transferred to offspring of that pregnancy.

Studies in humans may shed light on the properties of antibody populations that may change affecting efficiency of placental transport. Human maternal antibodies transferred to the fetus are IgG with a strong preference for IgG1(33-37). Furthermore, there is evidence for subpopulations of IgG1 that are preferentially transferred (38). Antibodies transferred to the fetus contain preferentially an Fc domain that efficiently binds to FcRn (Fc receptor neonate), which is largely responsible for transplacental transit of antibodies (33, 39, 40). In addition, it is reported that antibodies transferred are preferentially those with NK cell activating activity and are preferentially modified with galactose containing oligosaccharide side chains (41, 42). How these properties of maternal antibodies may evolve with time after immunization (or infection), pregnancies, or with maternal age is not clear. Notably, it has been reported that galactose content as well as sialic acid content of the Fc domain of antibodies decreases with age (43) but the mechanisms involved are not well understood. Results presented here, the differential levels of protection transferred to the cotton rat pups in breed 1 and breed 2, are consistent with such an evolution in properties of antibodies in immunized dams to those less compatible with placental transfer. It will be important in considerations of the use of maternal RSV vaccines to determine how a second immunization (boost) of CR dams during a second pregnancy affects the efficiency of transfer of protective antibodies to offspring of that pregnancy.

## Methods

### Preparation, characterization, and validation of VLP stocks

VLPs used as immunogens were based on the core proteins of Newcastle disease virus (NDV) M and NP proteins and contained the RSV F and G glycoproteins (44-46). The RSV proteins were assembled into the VLPs as chimera proteins with the sequences of the ectodomain of RSV F and G glycoproteins fused to the transmembrane and cytoplasmic domains of the NDV F and HN proteins, respectively. Three different VLPs were prepared, each containing the same RSV G chimera protein but with a different mutant F chimera protein. One VLP contained the DS-Cav1 pre-fusion F protein (30), while another VLP contained UC-3 F (30), a pre-fusion F protein with the cleavage site and intervening p27 sequences replaced with a seven-amino acid GS rich linker sequence as well as three point mutations, N67I, S215P, D486N, similar to the SC-TM F protein described by Krarup, et al. (47) A third VLP contained the post-F protein (30) and was used as a control, as previously described. The two pre-fusion F proteins also contained the foldon sequence inserted between the RSV F protein ectodomain and the NDV F protein transmembrane domain to stabilize further the pre-fusion conformation.

The VLPs were prepared by transfecting avian cells (ELL-0 from American Type Culture Collection) with cDNAs encoding the NDV M and NP proteins, the G protein chimera, and one of the mutant F chimera proteins (DS Cav1, UC-3 F, or post F). VLPs released into the cell supernatant were purified as previously described (48) and the F and G protein content of purified VLPs were quantified by Western blots and by monoclonal antibody (mAb) binding to the VLPs as previously described (18, 30). VLP stocks were adjusted for equivalent levels of F protein (18, 30). The pre-fusion or post fusion conformation of the F protein in the VLPs was validated by assessing the binding of mAbs specific to the pre-fusion form of the F protein to the VLPs as previously reported (18, 30).

### Preparation of soluble F proteins

Expi293F cells were transfected with cDNAs encoding the soluble DS-Cav1 pre-F protein or the soluble UC-3 pre-F protein. At six days post transfection, total cell supernatants were collected, cell debris removed by centrifugation, and the soluble polypeptides were purified on columns using the His tag and then the streptavidin tag (18, 49). Purified soluble DS Cav1 pre-F protein and soluble UC-3 pre-F protein efficiently bound to pre-fusion specific mAbs AM14 and D25 (30).

### Quantification of NP, M, H/G and VLP associated F proteins or soluble F proteins

For Western blots, proteins were resolved on 8% Bis-Tris gels (NuPage, ThermoFisher/Invitrogen). Quantifications of NP, M, H/G proteins, and RSV F/F proteins in VLPs or in soluble F protein preparations (pre-F, post-F) were accomplished after their separation in polyacrylamide gels followed by silver staining (Pierce Silver Stain, ThermoFisher) or Western blots of the proteins in parallel with protein standards as previously described (18, 49).

### ELISA

For determination of anti-pre F protein IgG antibody titers, wells of microtiter plates (ThermoFisher/Costar) were coated with either purified soluble DS Cav1 F protein or soluble UC-3 F protein (30 ng/well) and incubated overnight at 4°C, then blocked with 2% BSA for 16 hours. Different dilutions of sera, in PBS-2% BSA and 0.05% Tween, were added to each well and incubated for 2 hours at room temperature. Wells were then washed with PBS, incubated with chicken anti-cotton rat IgG antibody (Abnova PAB29753) coupled to HRP, and incubated for 1.5 hours at room temperature. Bound HRP was detected using TMB (3,3’5,5’-tetramethylbenzidin, ThermoFisher34028) and the reaction was stopped with 2N sulfuric acid. Color was read in SpectraMax Plus Plate Reader (Molecular Devices) using SoftMax Pro software. Amounts of IgG (ng/ml) in each dilution were calculated using a standard curve generated using defined amounts of purified cotton rat IgG.

### RSV Neutralization

RSV was grown in Hep2 cells, and RSV plaque assays were accomplished on Hep2 cells as previously described (18, 49). Antibody neutralization assays in a plaque reduction assay have been previously described (49). Neutralization titer was defined as log_2_ of the reciprocal of the dilution of serum that reduced virus titer by 60%.

### Animals

*Sigmodon hispidus* cotton rats (CR) were obtained from the inbred colony maintained at Sigmovir Biosystems, Inc (Rockville, MD). All studies were conducted under applicable laws and guidelines and after approval from the Sigmovir Biosystems, Inc. Institutional Animal Care and Use Committee. Animals were housed in large polycarbonate cages and fed a standard diet of rodent chow and water ad libitum. Animals were pre-bled before inclusion in the study to rule out the possibility of pre-existing antibodies against RSV. The colony was monitored for antibodies to paramyxoviruses and rodent viruses and no such antibodies were found. All cotton rats born as a result of breeding during these studies were used for RSV challenge at 4, 6, or 8 weeks of age, as indicated, and are referred to as “pups” or “offspring”.

### Quantification of bone marrow associated anti-pre-F IgG secreting long lived plasma cells (LLPC)

CR bone marrow cells were prepared as previously described (31). To quantify LLPC in bone marrows, wells of ELISpot plates (Millipore) were coated overnight with purified soluble pre-F protein (30 ng/well in PBS). Wells were washed and blocked for one hour in compete media. Four-fold serial dilutions of bone marrow cells were added, in triplicate to pre-coated wells and incubated at 37°C for 6 hours. Plates were washed and blocked overnight in PBS containing 1% BSA. Wells were incubated with HRP conjugated chicken anti-cotton rat IgG (1/2000 dilution of Abnova PAB29753) in PBS containing 1% BSA, incubated at room temperature, washed, incubated AEC substrate (BD Biosciences AEC substrate set, 551951) at room temperature until spots appear. Wells were washed and spots counted using CTL Immunospot S5 (46, 50)

### Experimental design

Three-week-old female cotton rats (dams) were tagged and separated into the indicated groups (Figure 1, panel B). Female cotton rats were bled and then the indicated groups were primed by RSV A/Long intranasal infection using a dose of 10^5^ PFU/animal in 50 μl. Eight weeks later (d56), females were paired with RSV negative males ∼2 weeks older than the females for mating (breed 1). Females were bled for serum collection at days 70, 84 (just before delivery), 141, 184, 231, 277, and 318 post priming. At day 70 different groups of primed and pregnant cotton rats were immunized with UC-3 F VLPs with 25, 75, 100, or 150 μg total VLP protein/animal (5, 15, 20, or 30 μg F protein), or mock immunized with PBS buffer (Figure 1). Other groups of pregnant cotton rats were immunized with DS-Cav1 F VLPs or post-F VLPs with 100 μg total VLP protein/animal (20 μg F protein) (Figure 1). Dams delivered pups at approximately day 84. Dams were bred again at day 156 without additional immunization (breed 2). Breed 2 pups were delivered on or about day 184. Breed 1 pups were bled and challenged with RSV A/Long (10^5^ PFU/animal) at 4 or 6 weeks of age. Breed 2 pups were bled and challenged with RSV at 4 or 8 weeks after birth. All pups were sacrificed on day 4 post challenge. Pup serum NAb and total anti-pre-F IgG and nose and lung viral titers were measured, as previously described (18). Dams were kept in the study for an additional 134 days after delivery of breed 2 pups for additional serum collection.

### Statistical analysis

Statistical analyses (student-*t* test) of data were accomplished using Graph Pad Prism 9 software.

## Acknowledgements

This research was funded by grants AI109926 (J.C.G.B) and AI043896 (TM) from the National Institutes of Health. We thank Jason McLellan and Judy Beeler for F protein mAbs. We thank Steven Hatfield for significant technical help. We thank Mr. Charles Smith, Martha Malache, and Ana Rivera for the animal care service.

